# Emergence and rapid global dissemination of CTX-M-15-associated *Klebsiella pneumoniae* strain ST307

**DOI:** 10.1101/352740

**Authors:** Kelly L. Wyres, Jane Hawkey, Marit A.K. Hetland, Aasmund Fostervold, Ryan R. Wick, Louise M. Judd, Mohammad Hamidian, Benjamin P. Howden, Iren H. Löhr, Kathryn E. Holt

## Abstract

Recent reports indicate the emergence of a new carbapenemase producing *Klebsiella pneumoniae* clone, ST307. Here we show that ST307 emerged in the mid-1990s (nearly 20 years prior to its first report), is already globally distributed and is intimately associated with a conserved plasmid harbouring the *bla*_CTX-M-15_ extended-spectrum beta-lactamase (ESBL) gene plus other antimicrobial resistance determinants. Our findings support the need for enhanced surveillance of this widespread ESBL clone in which carbapenem resistance is now emerging.

## Background

Several reports have indicated the recent emergence of a new multi-drug resistant (MDR) *Klebsiella pneumoniae* (*Kp*) clone, ST307. We have recently generated data suggesting that ST307 is becoming an important cause of extended-spectrum beta-lactamase (ESBL) producing *Kp* infections in Norway (unpublished), and others have reported it as an emerging cause of carbapenemase-producing *Kp* infections [1–3].

Here we summarise the available literature and investigate 95 ST307 whole-genome sequences to better understand the emergence and molecular epidemiology of this clone and identify the antimicrobial resistance (AMR) genes with which it is associated.

## Growing reports of ST307 infections

The oldest recorded ST307 isolate was collected in The Netherlands in 2008 (*Kp* MLST database; bigsdb.pasteur.fr/klebsiella/klebsiella.html). The earliest clinical strain reported in the literature was collected in Pakistan in 2009 [4] and was followed by sporadic isolations across Europe, Asia, Africa and the Americas (summarised in **Table S1** and **Figure 1**). Several reports have indicated local dissemination of ST307 harbouring *K. pneumoniae* carbapenemase (KPC) genes, *bla*_KPc-2_ or *bla*_KPC-3_ [3,5,6], while an analysis of >1700 ESBL-producing *Kp* from a hospital network in Texas, USA found high prevalence of *bla*_CTX-M-15_-postive ST307 strains, ~1/3 of which also carried *bla*_KPC_ genes [1]. This was consistent with other reports that bla_CTX-M-15_ is common in ST307 [1–4,7]. Aside from *bla*_KPC_ genes, carbapenem resistance conferred by NDM-1 or OXA-48 carbapenemases has been reported [1–3,7,8], as has resistance to the novel beta-lactam-inhibitor combination, ceftazidime-avibactam [9], and colistin [8,10].

**Figure 1:**
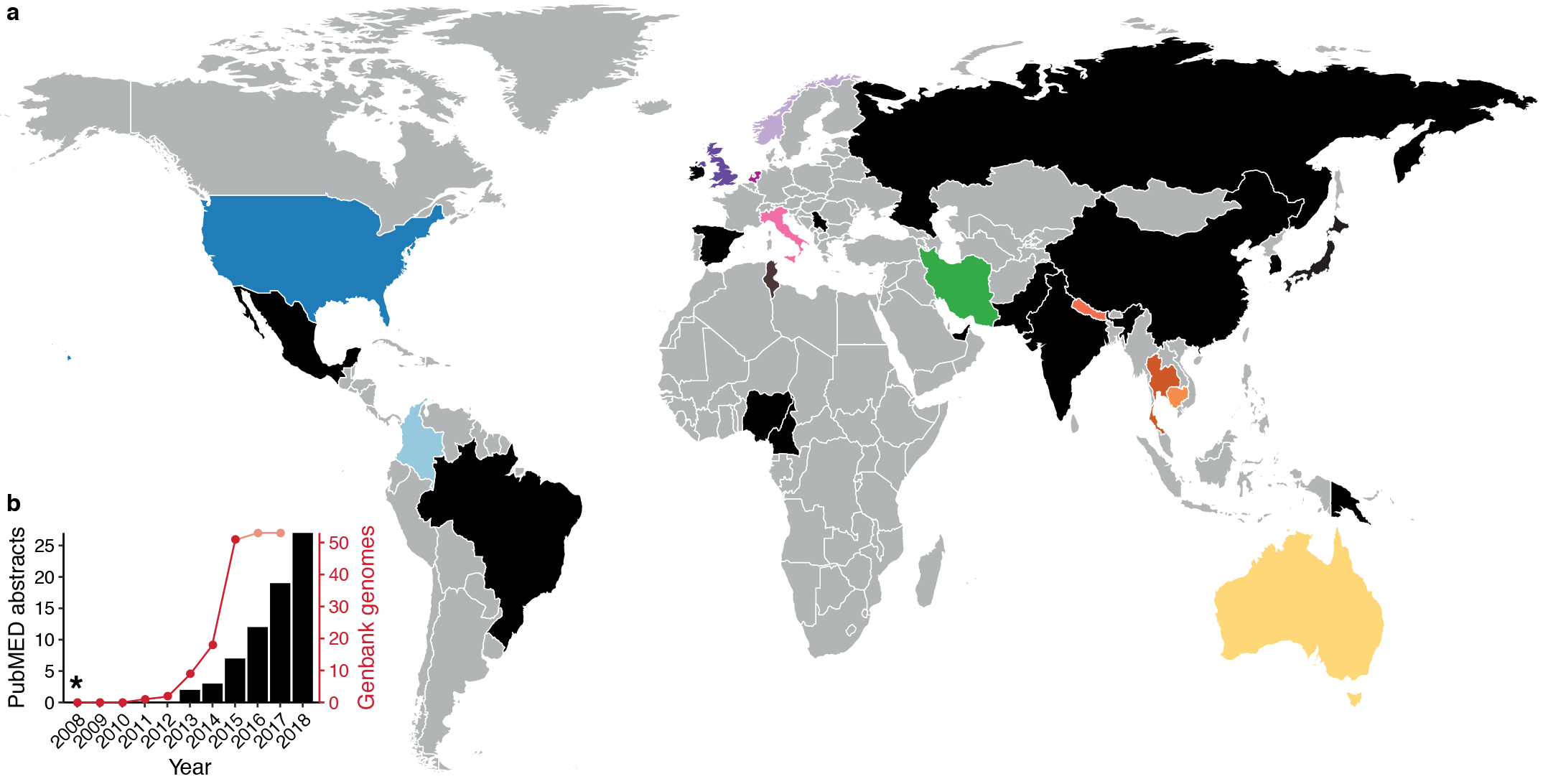
Geographic distribution and increasing reports of ST307. **a,** Countries of collection of ST307 isolates reported in the literature, international *K. pneumoniae* MLST database and/or for which genome data are available (also see **Table S1**). Countries for which corresponding isolate genomes were available for the current analysis are coloured as per **Figure 2**. All other countries from which ST307 has been reported are coloured black. **b,** Reports of ST307 in the literature and among genome assemblies deposited in Genbank. Black bars show the cumulative number of PubMED abstracts as of April 2018, identified using the search criteria “ST307” with/without “*Klebsiella pneumoniae*,” also see **Table S1**. Red/peach line shows the cumulative number of isolates for which genome assemblies are deposited in Genbank as of Dec 2017; note dates indicate year of isolate collection not date of deposition, hence peach line indicates values that will likely increase as further genomes are deposited. *indicates the date of collection of the first ST307 isolate reported in the international MLST database.

## ST307 genome data and analysis

We collected 504 published and 37 newly sequenced ST307 genomes representing 11 countries on five continents. For the latter, 150 or 250 bp PE reads were generated on the Illumina MiSeq platform as described previously [11]. Norwegian isolate collection was approved by REC West (Norway) ethics committee (application ID: 2017/1185). The oldest isolate (Kp616 from Iran, 2009) was also subjected to long read Oxford Nanopore sequencing and hybrid genome assembly using *Unicycler* v0.4.4 [12,13]. The completed Kp616 genome comprised a 5,246,307 bp chromosome plus two plasmids (pKp616_1, 58 kbp; pKp616_2, 55 kbp; GenBank accession GCA_003076555.1). All sample information, accession numbers and citations are listed in **Table S2**.

A core chromosomal single nucleotide variant (SNV) alignment was generated using *RedDog* (reference: Kp616 chromosome) as described in [11]. Recombination was removed using *Gubbins* [14]. A preliminary tree using *FastTree* [15] indicated the majority of strains from Texas (n=451/468, 96.4%) formed a distinct monophyletic clade, hence we randomly selected one isolate per year to represent this clade in comparative analyses. The final recombination-free alignment of 1,465 SNVs in 95 genomes was subjected to Bayesian phylogenetic analysis using *BEAST2* v2.4.7 [16] as described in [11] (**Figure 2**). A GTR, relaxed clock, constant population size model was determined to be the best fit and we confirmed a strong temporal signal by date-randomisation and linear regression (**Figure S1**).

**Figure 2:**
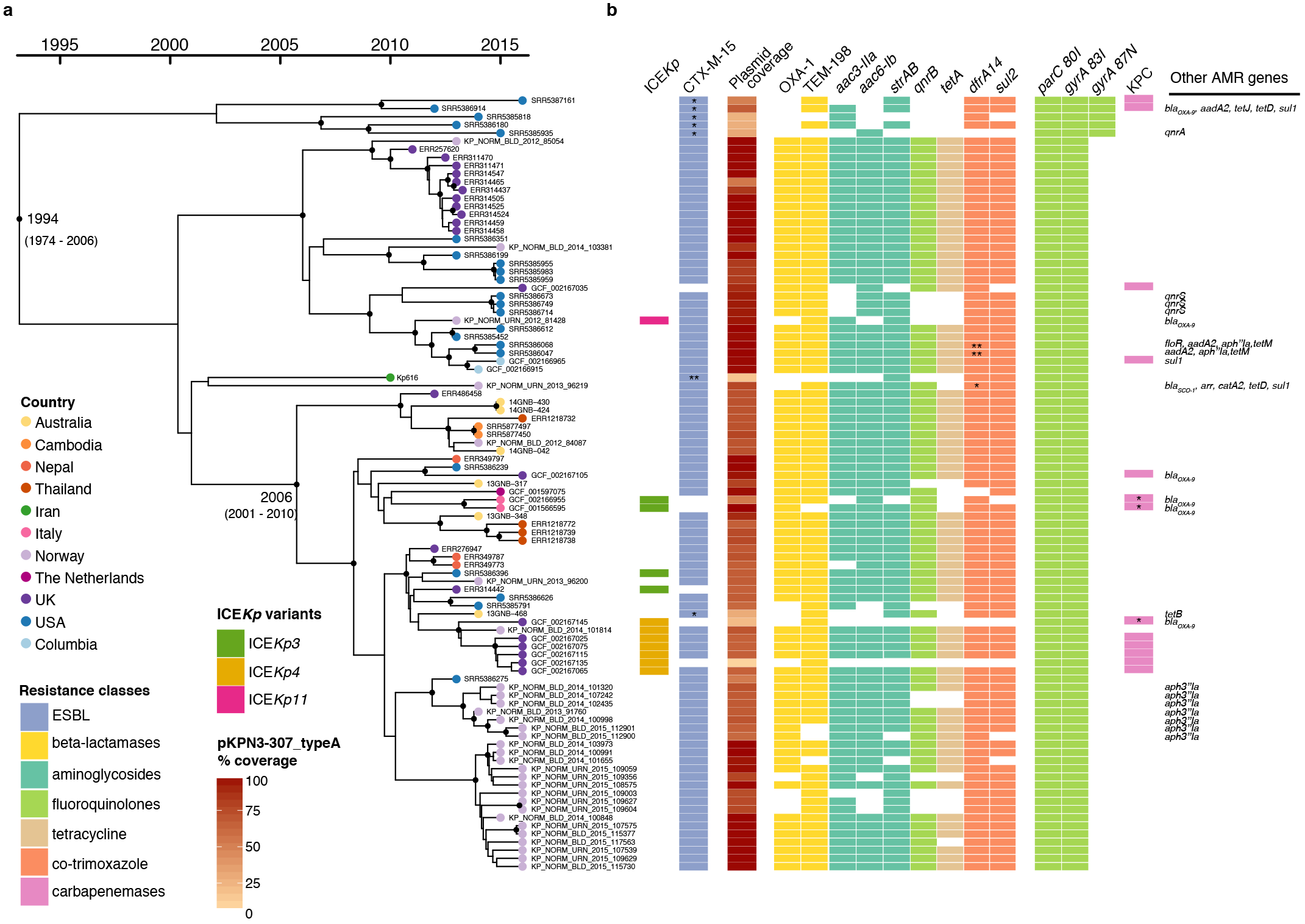
Bayesian phylogeny of 95 ST307 isolates. **a,** Dated phylogeny of ST307 isolates (n=95), with tips coloured by country of isolation (as per inset legend and **Figure 1**). Black dots on internal nodes indicate ≥95% posterior probability. **b,** Presence of the yersiniabactin-carrying ICE*Kp* elements (variants coloured as per inset legend), antimicrobial resistance genes (blocks coloured by drug class), and of the *bla*_CTX-M-15_ plasmid pKPN3-307_typeA. In the CTX-M-15 column, * indicates *bla*_CTX-M-15_ is inserted in chromosome, ** *bla*_CTX-M-15_ inserted in IncN plasmid; in the *dfrA14* column, * *dfrA27* allele, ** *dfrA12* allele; in the KPC column, * indicates *bla*_KPC-3_, otherwise *bla*_KPC-2_ is present.

## ST307 emerged in the mid-1990s and has disseminated globally

We estimate that ST307 emerged in 1994 (95% HPD, 1974-2006), close to the emergence date estimated previously for ST258 [17], despite the fact that the latter was reported in the literature and recognised as a disseminated clone more than a decade earlier than ST307 [18]. The estimated mutation rate for ST307 (1.18×10^−6^ substitutions site^−1^ year^−1^, 95% HPD, 8.01×10^−7^-1.58×10^−6^; **Figure S1**) was also remarkably similar to that estimated previously for ST258 [17].

The phylogeny revealed two deep-branching lineages, one of which has become globally distributed, comprising genomes from the Americas, Asia, Australia, the Middle East and Europe (**Figure 2**). Within this lineage there was evidence of transfer of ST307 between countries, and for all countries with >3 genomes there were multiple clusters within the global lineage. The countries with the highest representation were distributed most broadly (Norway, n=30; USA, n=22; UK, n=22), suggesting that the same patterns would likely be detected for most countries if sampling was increased (**Figure 2**). The second lineage included only subsampled strains from Texas (2011-2015), indicating that the majority of the hundreds of infections attributed to ST307 in [1] resulted from prolonged local transmission of this clade. Texan isolates were also found in the global lineage, suggesting that the USA may be a potential origin for ST307, as most of its genetic diversity was present in that location.

## ST307 virulence and antimicrobial resistance genes

We used *Kleborate* [19] to investigate the prevalence of acquired AMR and virulence determinants and *Kaptive* [20] to investigate capsule (K) and lipopolysaccharide (O) locus diversity. We detected no evidence of the *Kp* virulence plasmid [21], but a minority of genomes (n=12, 12.6%) harboured the yersiniabactin siderophore locus located within three distinct chromosomally integrated ICE*Kp* variants. The distribution of ICE*Kp* insertions on the ST307 core genome tree indicated >4 independent acquisitions with limited expansion of recipient sublineages (**Figure 2**), consistent with the patterns recently reported for ST258 and other common MDR *Kp* clones [19]. Unlike those other clones [17,22], all ST307 shared the same capsule (KL107) and O antigen (O2v2) loci. An additional putative capsule synthesis locus (Kp616 genes C2861_20465 to C2861_20520 [2]), which is unlike any known K-locus, was also conserved among ST307 but is rare among the broader *Kp* population (data not shown).

In contrast to the virulence loci, acquired AMR genes were highly prevalent; 93 (97.9%) isolates carried acquired resistance determinants associated with ≥3 drug classes (**Table S2**). The ParC 80I and GyrA 83I fluoroquinolone resistance-associated mutations were conserved in all genomes. The *bla*_CTX-M-15_ ESBL gene was found in 89 (93.7%) genomes, and 81 (85.3%) harboured it in combination with *sul2*, *dfrA14* and *strAB* with/without *aac(3)-IIa*, which were all linked to an MDR plasmid (see below). *bla*_KPCs_ and other AMR genes were occasionally identified (see **Figure 2** and **Table S2**).

For the majority of genomes carrying *bla*_CTX-M-15_ (n=88/89) BLASTn confirmed that this gene was located downstream of *ISEcp1*, which forms a transposon to mobilise *bla*_CTX-M-15_ and promotes its expression [23]. Four complete ST307 IncFII_K_/IncFIB_K_ *bla*_CTX-M-15_ plasmids have been published [2], and share an insertion of the IS*Ecp1/bla*_CTX-M-15_ transposon within Tn3. Read-mapping to pKPN3-307_typeA (accession KY271404) and assembly graph inspections of our *bla*_CTX-M-15_-positive genomes showed that all carried the same IS*Ecp1*/*bla*_CTX-M-15_ transposon. In 84/89 cases, the same pKPN3-307_typeA plasmid backbone was present and IS*Ecp1*/bla_CTX-M-15_ was located in the same site within Tn3, consistent with conservation of the same ESBL plasmid (**Figure S2**). The exceptions were: (i) an Australian isolate (13GNB-468) had the IS*Ecpl*/*bla*_CTX-M-15_ transposon inserted in the chromosomal gene *feoB*; (ii) Kp616 carried no pKPN3-307_typeA-like plasmid but harboured *bla*_CTX-M-15_ on an IncN plasmid; and (iii) the five representatives of the Texan-specific lineage carried two chromosomal insertions of the IS*Ecp1*/*bla*_CTX-M-15_ transposon (within Kp616 loci C2861_02545 and C2861_22795). This coupled with the additional GyrA 87N fluoroquinolone resistance mutation in the Texan lineage may have contributed to its prolonged transmission in the hospital setting. Regardless of these exceptions, the level of plasmid conservation in ST307 is remarkable, mirroring the association of ST258 with the *bla*_KPC_ pKpQIL plasmid [17] and suggesting that the plasmid confers limited fitness cost to the host (although plasmid-positive, *bla*_CTX-M-15_-negative genomes were observed, **Table S2**).

## Conclusions

Complementing the increasing reports in the literature, our analyses reveal that ST307 is a highly successful MDR clone that shares many traits with ST258, but is closely associated with *bla*_CTX-M-15_ rather than *bla*_KPCs_. With sufficient exposure ST307 can acquire and disseminate carbapenemases [1–3], and likely other clinically important AMR determinants. ST307 appears to be readily transferred between countries and has already become globally disseminated, yet has remained largely unnoticed for almost 20 years. These findings indicate an urgent need for enhanced surveillance of MDR *Kp* to monitor ST307 alongside other well-known clones and detect emerging MDR threats.

## Acknowledgements

This work was supported by the Viertel Foundation of Australia, the NHMRC of Australia (fellowship #GNT1105905 to BPH), The Western Norway Regional Health Authority (fellowship 912037 and 912119, and grant number 912050) and the Bill and Melinda Gates Foundation, Seattle. We thank; The Norwegian Surveillance System for Antimicrobial Drug Resistance (NORM) for data sharing; The Norwegian *Klebsiella pneumoniae* study group for collection of Norwegian isolates; Dr Mohammad Ali Boroumand for collection of the Iranian isolate; the Australian Group on Antimicrobial Resistance (AGAR) in particular Jan Bell, for providing isolates from Australia, and the staff at MDU Public Health Laboratory for sequencing Australian isolates.

## Author contributions

KLW, JH, IHL and KEH conceived the study and performed data analyses. MAKH performed systematic literature review. AF, IHL, MH and BPH provided isolates and/or genome data. RRW and LMJ generated the completed reference genome. KLW, JH and KEH wrote the paper. All authors contributed to data interpretation, read and commented on the manuscript.

